# A novel and accurate full-length HTT mouse model for Huntington’s disease

**DOI:** 10.1101/2021.06.14.448318

**Authors:** Sushila A Shenoy, Sushuang Zheng, Wencheng Liu, Yuanyi Dai, Yuanxiu Liu, Zhipeng Hou, Susumu Mori, Wenzhen Duan, Chenjian Li

## Abstract

Here we report the generation and characterization of a novel Huntington’s disease (HD) mouse model BAC226Q by using a bacterial artificial chromosome (BAC) system, expressing full-length human HTT with ∼226 CAG-CAA repeats and containing endogenous human HTT promoter and regulatory elements. BAC226Q recapitulated a full-spectrum of age-dependent and progressive HD-like phenotypes without unwanted and erroneous phenotypes. BAC226Q mice developed normally, and gradually exhibited HD-like mood and cognitive phenotypes at 2 months. From 3-4 months, BAC226Q mice showed robust progressive motor deficits. At 11 months, BAC226Q mice showed significant reduced life span, gradual weight loss and exhibit neuropathology including significant brain atrophy specific to striatum and cortex, striatal neuronal death, widespread huntingtin inclusions and reactive pathology. Therefore, the novel BAC226Q mouse accurately recapitulating robust, age-dependent, progressive HD-like phenotypes will be a valuable tool for studying disease mechanisms, identifying biomarkers and testing gene-targeting therapeutic approaches for HD.

## Introduction

Huntington’s disease (HD) is an autosomal-dominant hereditary neurodegenerative disorder caused by a pathogenic expansion of the CAG trinucleotide repeats in exon 1 of the huntingtin (HTT) gene (The Huntington’s Disease Collaborative Research Group, 1993). The clinical features of HD include motor deficit, psychiatric disturbance, cognitive impairment, and peripheral signs such as weight loss and sleep disturbance (Ghosh & Tabrizi, 2015). Disease onset, which is dependent on the CAG repeat length, is usually defined by the onset of motor symptoms, although non-motor symptoms are often present many years in advance. The striking neuropathological characteristic of HD, induced by the mutant Htt protein (mHtt), is the progressive brain atrophy mainly in the striatum and cerebral cortex (Walker, 2007). Despite ubiquitously expressed mHtt protein in HD brain, the primary vulnerable neuronal types are medium spiny neurons (MSNs) in the striatum and pyramidal neurons in the cortex (Vonsattel & DiFiglia, 1998). Widespread mHtt protein aggregation in HD brain tissue is another hallmark of HD pathology (Davies & Scherzinger, 1997). While the causative gene for HD was identified two decades ago, there are still no effective disease-modifying therapies available to prevent or delay the progression of this disorder.

Since the identification of causative gene for HD, a large number of animal models have been generated in a variety of animal species including *C. elegans, Drosophila melanogaster,* zebra fish, mouse, rat, sheep, pig and non-human primates to better elucidate the complex pathogenic mechanisms in HD, and to develop potential therapeutic strategies for HD (Farshim & Bates, 2018). Although large animal models such as pig and non-human primates are very useful to study HD (Niu *et al*., 2010; Sasaki *et al*., 2009; Yan *et al*., 2018; Yang *et al*., 2008), mouse models still dominate the research field and provide us with valuable tools to investigate the pathogenesis of HD and therapeutic targets.

To date, there are more than 20 different HD mouse models available (Brooks & Dunnett, 2015; Crook & Housman, 2011; Farshim & Bates, 2018; Pouladi, Morton, & Hayden, 2013). In general, those genetic mouse models can be classified into three groups based on distinct strategies: the transgenic models carrying human HTT N-terminal fragments with CAG expansions such as R6/1, R6/2 and N171-82Q (Carter *et al*., 1999; Mangiarini *et al*., 1996; Schilling *et al*., 1999), the transgenic full-length mutant HTT models such as YAC128, BACHD, BAC-225Q (Gray *et al*., 2008; Slow *et al*., 2003; Wegrzynowicz *et al*., 2015) and humanized HD mice (Hu97/18 and Hu128/21) (Southwell, Skotte, Villanueva, Ostergaard, *et al*., 2017; Southwell *et al*., 2013), and knock-in models with modified CAG repeats length of endogenous mouse HTT gene such as HdhQ72, HdhQ94, HdhQ111, HdhQ140, HdhQ150, zQ175, N160Q and Q175FDN (Heng, Tallaksen-Greene, Detloff, & Albin, 2007; Hickey *et al*., 2008; Levine *et al*., 1999; Lin *et al*., 2001; Liu *et al*., 2016; Menalled *et al*., 2012; Menalled, Sison, Dragatsis, Zeitlin, & Chesselet, 2003; Shelbourne *et al*., 1999; Southwell *et al*., 2016; Wheeler *et al*., 1999). The existing mouse models mimic some aspects of HD including behavioral disturbances and neuropathological changes of the disease. However, none of them can fully recapitulate human disease, and many of them have spurious phenotypes that are irrelevant or opposite to human conditions. The N-terminal fragment transgenic mice such as R6/2 show a robust phenotype and severely reduced life span, but they demonstrated severe developmental deficits and nonspecific neurodegeneration throughout the central nervous system (Carter *et al*., 1999; Crook & Housman, 2011; Heng *et al*., 2007; Mangiarini *et al*., 1996; Southwell *et al*., 2016) as well as diabetes, which are not correlated with HD patients. In comparison, full-length transgenic HD mice and Hdh knock-in mice replicate relatively better the neuropathology of HD, however they display much milder phenotypes and slower progression (Brooks, Jones, & Dunnett, 2012; Farshim & Bates, 2018; Pouladi *et al*., 2013). Moreover, YAC128, BACHD, and humanized HD strains (Hu97/18 and Hu128/21) exhibit significant weight gain, opposite to what is seen in HD patients (Gray *et al*., 2008; Slow *et al*., 2003; Southwell, Skotte, Villanueva, Ostergaard, *et al*., 2017; Southwell *et al*., 2013). Another commonly used mouse model zQ175 KI has robust phenotypes (Lin *et al*., 2001), however it has mouse HTT gene and therefore not suitable for testing the approach of genetically deleting human mutant HTT gene via methods such as CRISPR-Cas9.

Therefore, despite all the achievements, we are still in search for a new full-length model that not only accurately recapitulates all HD phenotypes, but also without spurious characteristics. In light of the clear relationship between PolyQ repeat length and disease onset and severity in both humans and mice, in this study, we engineered a bacterial artificial chromosome (BAC) transgenic mouse model of HD-BAC226Q which expresses full-length human HTT with 226Q encoded by a mixture of CAG-CAA repeat. This 226Q repeat length is smaller than the maximum 250 polyQ in HD patient cases (Nance, Mathias-Hagen, Breningstall, Wick, & McGlennen, 1999). The CAG-CAA mix will stabilize the polyQ length. Our novel BAC226Q mouse recapitulates a full spectrum of cardinal HD symptoms and pathologies: reduced life span, weight loss, motor and non-motor neurologic phenotypes, selective brain atrophy, striatal neuronal death, mHtt aggregation and gliosis. It is also important to note that there are no spurious abnormalities identified to date. Therefore, the new HD mouse model will be valuable for studying pathogenic mechanisms and developing therapeutics.

## Results

### Generation of BAC226Q transgenic mice

Transgenic BAC226Q mice were generated and contain the full-length human HTT with 226 CAG-CAA repeats under the control of the endogenous human HTT promoter and regulatory elements. Similar to the BACHD mouse, a mixed CAG-CAA repeat is used to create a stable length of polyglutamine (PolyQ) that is not susceptible to expansion or retraction (Gray *et al*., 2008). To determine the copy number and insertion sites in FVB mouse genome of full-length human HTT, whole genome sequencing was applied and indicated two copies of the human HTT were inserted in the chromosome 8 in FVB mouse genome at Chr8:46084002 (Fig 1A). The protein expression level of HTT was determined by Western blots with whole brain lysates from 2- and 11-month old animals probed with anti-polyQ MAB1574 (1C2 clone) and S830 antibody against mutant huntingtin protein (Fig 1B and Fig 1C). Age-matched 11-month BACHD and wildtype mice were used for comparison. As expected, the band for 226Q mutant huntingtin in BAC226Q mouse is at a higher molecular weight than that for 97Q huntingtin in the BACHD mouse. There are no Htt protein fragments detected in 2-month or 11-month BAC226Q mouse in Western blots (Fig 1B).

**Figure 1.**
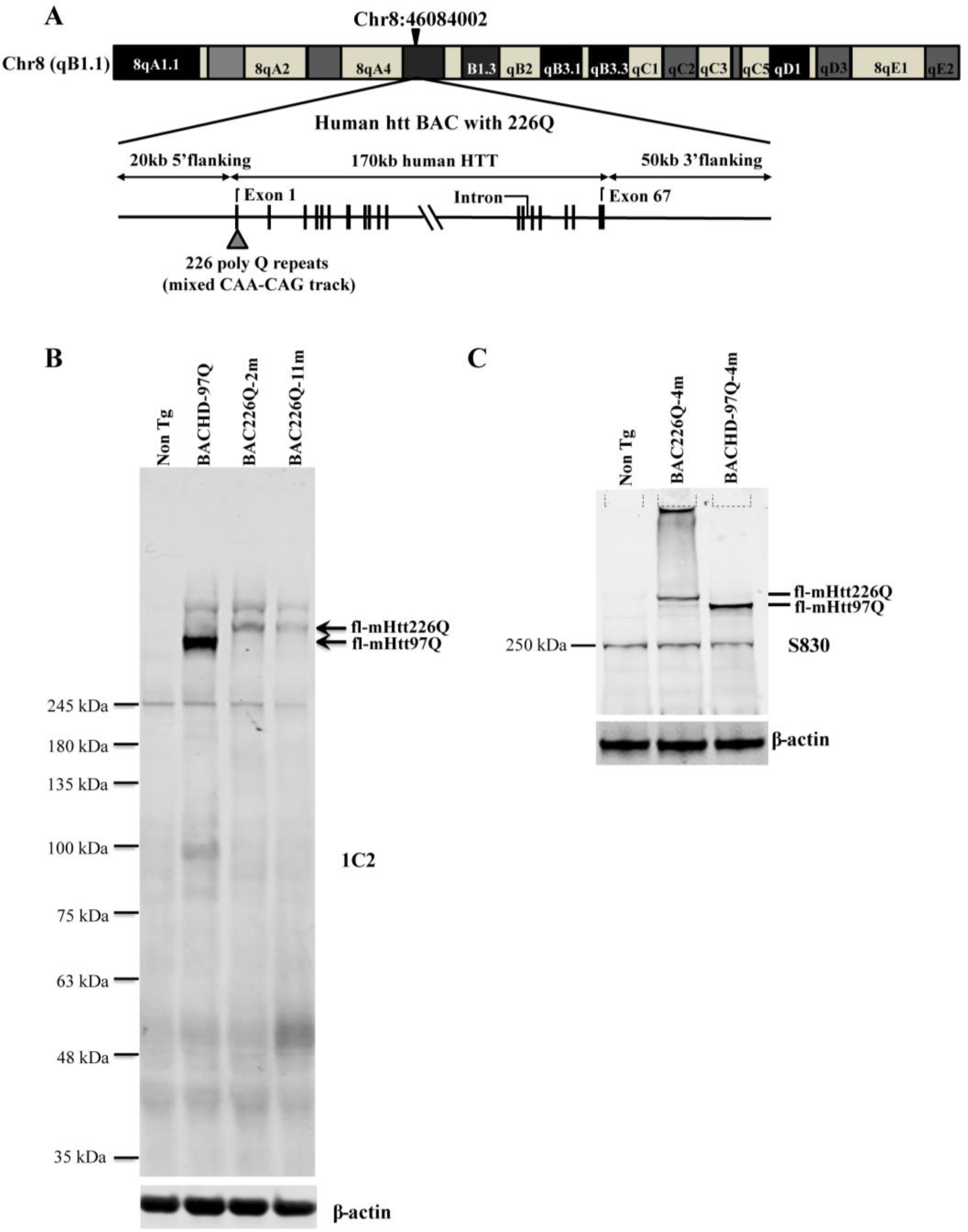
Generation of BAC226Q transgenic mice. A) Schematic diagram of the insertion of the full-length human mutant HTT BAC into mouse genome. The human HTT BAC contains 170kb genomic DNA of complete HTT genomic locus plus approximately 20 kb 5’ and 50 kb 3’ flanking region with endogenous regulatory elements. A mixed CAA-CAG repeat encoding 226 polyQ was engineered in the Exon 1. By whole genome sequencing, two copies of the human mHTT BAC had been detected and inserted into the mouse genome at Chr8:46084002. B) Western blot analysis of Htt protein expression levels in BAC226Q, non-transgenic littermate and BACHD (97Q) mice with antibody 1C2. The whole-brain lysates are from 2- and 11-month BAC226Q, 11-month non-transgenic littermate and BACHD mice. Western blots were repeated in 3 independent cohorts of mice. The upper panel: arrows indicate that 1C2, an antibody specific for mutant Htt, detects mHtt in BAC226Q and BACHD (97Q) mice but not non-transgenic littermate. In the second lane from the left, mHtt-97Q from BACHD (97Q) mouse appeared at the expected molecular weight. In the right two lanes, mHtt-226Q from 2 and 11-month BAC226Q mice are detected at a molecular weight higher than that of mHtt-97Q. Lower panel: the same blot was probed with anti-β-actin antibody for the loading control. C) Reconfirmation by Western blot with the S830 antibody specific for mHtt in 4-month non-transgenic control, BAC226Q and BACHD (97Q) mice. S830 antibody detected mHtt-97Q in BACHD (97Q) mouse (right lane) and mHtt-226Q (middle lane) at expected molecular weights but not in non-transgenic control (left lane).

### Progressive weight loss and shortened life span in BAC226Q mice

As a first step in characterizing these mice, their growth and reproduction were examined. Both male and female transgenic mice were born in the expected Mendelian ratios with no obvious abnormalities. Both males and females were fertile, but males were preferred for breeding because female animals had a short window of time during which they were able to adequately care for pups. Animal weights were recorded weekly and the results show that HD mice were born normal and gained weight at the same rate as their non-transgenic littermates in the first 8 weeks of development (Fig 2B). In both male and female WT littermates, body weight continuously increased until the 15 month of age, however, BAC226Q exhibited slower weight gain followed by weight loss (Fig 2C, D). At 6 months of age, we observed significant reductions in body weight in BAC226Q mice compared to WT littermates (Fig 2C), and more than 50% loss of BAC226Q at 15 months (Fig 2C & Fig 2D). Weight loss is a common and serious complication in HD patients, who fail to maintain weight even when caloric intake is increased (Aziz *et al*., 2008). BAC226Q mice recapitulated this aspect accurately. Transgenic BAC226Q mice had significantly reduced lifespan (Fig 2A). In both sexes, only 50% of transgenic animals survive past 1 year. The longest surviving transgenic mice were 15 months old when they had to be euthanized due to the end-point condition according to animal welfare requirements. In comparison, non-transgenic littermate controls appeared healthy at 15 months of age. These initial results suggest that BAC226Q mice have progressive weight loss and shortened lifespan, both of which correlate well with the human disease condition.

**Figure 2.**
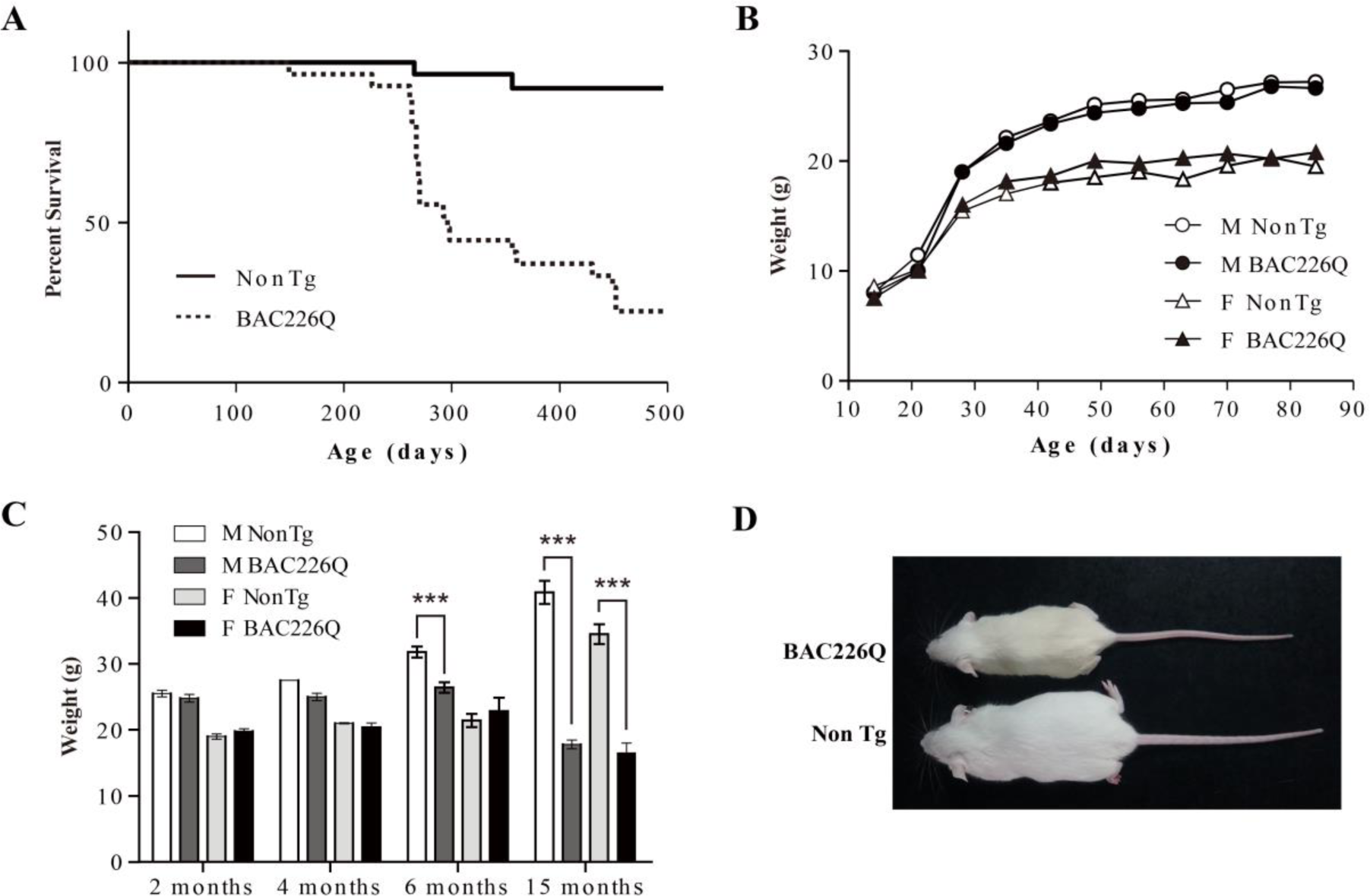
Shortened life span and weight loss in BAC226Q mice. A) Kaplan-Meier survival curves indicate a median life span of less than 1 year for BAC226Q mice (start with n=16-21). B) In the first 12 weeks, male and female BAC226Q mice gained weights at the same rate as their non-transgenic littermates (n=8-12). C) After normal development, beginning from 6 months, BAC226Q mice had progressive and significant weight loss compared to non-transgenic controls (n=8-12, ***=*p* <0.001). D) Representative body sizes of 11-month old male BAC226Q mouse and non-transgenic littermate.

### Age-dependent and progressive motor impairments in BAC226Q mice

The earliest, most visible motor phenotype in the BAC226Q mice was the chorea-like movement which first manifested between 12 and 14 weeks. The motor phenotypes include abrupt unnatural jerking and twisting of the head and body (Video 1), resembling the chorea movement in HD patients. It should be noted that every transgenic BAC226Q mouse exhibits this HD-like movement. Between 14 and 16 weeks, rapid circling behavior appeared, and was especially prominent when animals were disturbed in their cages (Video 2).

To characterize the progressive nature of motor deficits in our model, we measured motor function at serial time points by several behavioral tests. First, overall activity was measured in the open field task (Fig 3A, B). Open field activity is divided into horizontal and vertical components. Horizontal activity is measured by lateral movement around the open field, and vertical activity is measured by rearing movements. At 2 months, HD mice are indistinguishable from non-transgenic littermate controls in both horizontal and vertical activity measures. Horizontal activity is dramatically increased at 4 months (Fig 3A), but returns to wild-type levels at 10 and 15 months. In contrast, vertical activity of 4-month HD mice has declined to about 20% of the controls (Fig 3B). This suggests that in addition to their hyperkinesia, the mice cannot rear normally, indicating a loss of motor control. While reduced vertical activity at 4 months may be caused by the increase in horizontal movement, at 10 and 15 months, horizontal activity is reduced to wild-type levels but vertical activity does not recover and becomes even more impaired. These results parallel the biphasic motor symptom progression in HD patients in which early stage involuntary movements become more dystonic in late stages of disease.

**Figure 3.**
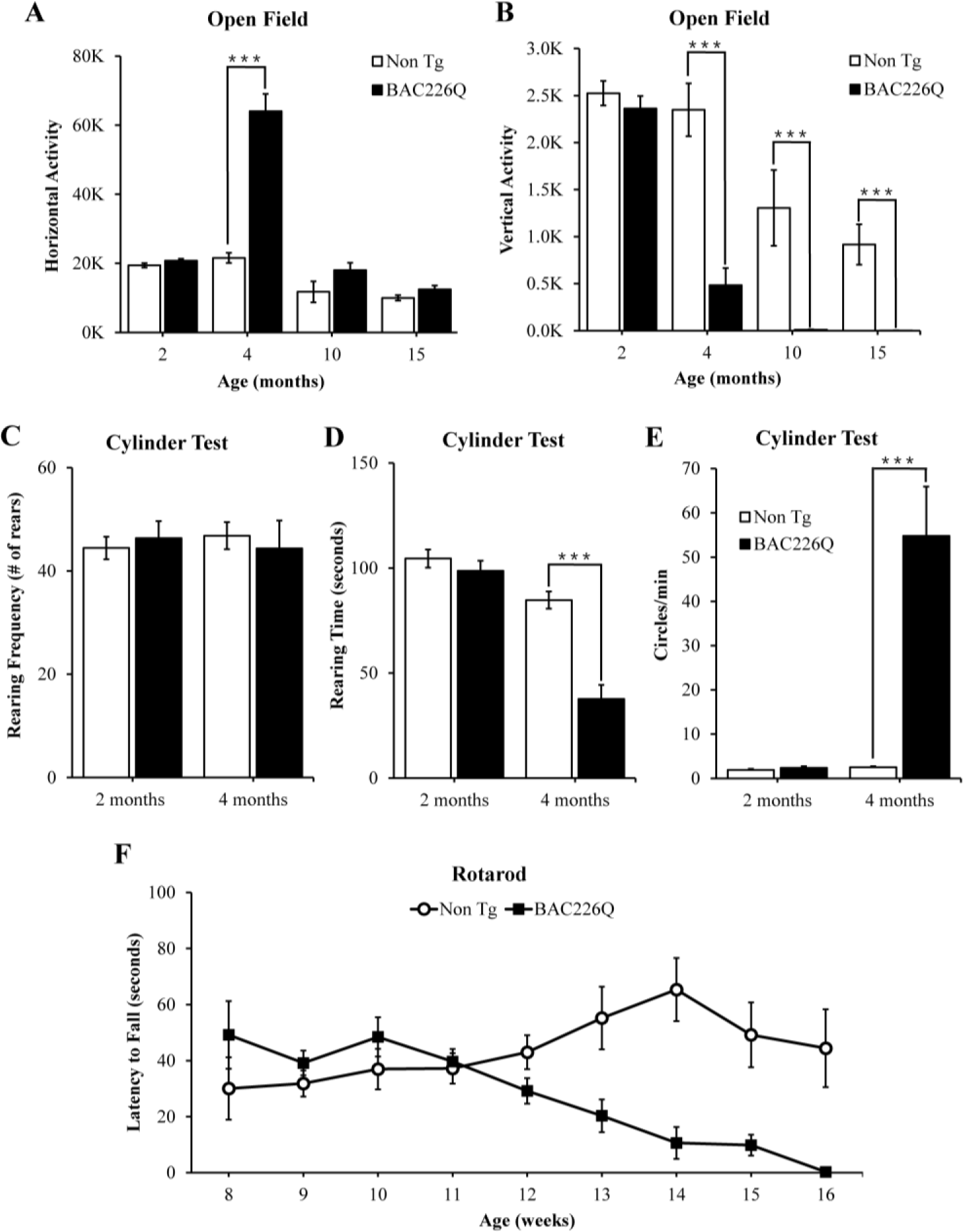
Robust, early onset and progressive motor deficits in BAC226Q mice. A & B) Horizontal and vertical activity are measured by total beam breaks in 1 hour in the open field. In horizontal movement, 4-month BAC226Q mice developed robust hyperactivity (A). In vertical movement, BAC226Q mice had progressively diminished activity (B). C, D, & E) Results of the cylinder task at 2 and 4 months. Rearing frequency, the total number of rears observed during the 5-minute task, is significantly reduced in BAC226Q mice (C). Rearing time, the total time of mice in an upright rearing position, is greatly reduced in BAC226Q mice (D). Circling frequency, the total number of clockwise and counterclockwise rotations, is obviously increased in BAC226Q mice (E). F) Rotarod performance was averaged over three trials, performed once a week. BAC226Q mice showed progressive and strong deficits from 12 weeks. In all tests, significance is indicated by ***= *p* <0.001, n = 11 per group.

The cylinder test was performed to provide another sensitive analysis of motor phenotypes. Similar to the open field test, BAC226Q mice are normal at 2 months and defective at 4 months in the cylinder test (Video 3). At 4 months, rearing frequency was unchanged, but rearing time was significantly reduced (Fig 3C, D). These results suggest that the transgenic mice retained their instinct to rear to explore the environment, but were unable to sustain upright posture due to motor impairment. The circling phenotype in BAC226Q mice (Figure 3E) was reminiscent of that seen in unilateral lesion models, but with important differences. In lesion models, stereotyped turning occurs in a single direction determined by which hemisphere is lesioned. While BAC226Q mice spent time turning in both directions, and the frequency of turning was significantly increased over controls. Circling in BAC226Q mice may be explained by a reduced capacity to inhibit or terminate movements such as turns.

To demonstrate deficits in balance and coordination and to track the progressive decline of motor function from normal at 2 months to severe at 4 months, an accelerating rotarod test was used. The accelerating rotarod requires mice to walk forward on a rotating rod to maintain balance and is a sensitive measurement of motor coordination. Consistent with the open field and cylinder results, no differences between BAC226Q and controls could be detected until 12 weeks (Fig 3F). At 13 weeks, BAC226Q mice performed significantly worse compared to non-transgenic controls. At 16 weeks, BAC226Q were unable to balance on the rotarod for any duration of time. These results demonstrated a rapidly progressing, debilitating motor dysfunction with a clearly defined time course.

### Cognitive and psychiatric-relevant deficits in BAC226Q mice prior to motor abnormalities

In HD patients, cognitive and psychiatric symptoms are common and as detrimental to quality of life as motor disorders. To characterize the non-motor phenotypes in the BAC226Q mice, we subjected 2-month-old BAC226Q mice to several tasks. Testing was performed at 2 months for two reasons: First, these tests depend to some degree on the ability of the animal to move around, thus we need to test at 2 months when there is no measurable motor impairment. Second, psychiatric and cognitive symptoms frequently occur years before the onset of motor symptoms in HD patients. The tests were performed in an ascending order of stress level, from object-in-place memory test, sucrose preference to the most stressful forced swim task.

Because there is an evidence of impaired hippocampal function in pre-symptomatic HD patients, we chose a challenging cognitive task with a hippocampus-dependent spatial component (Phillips, Shannon, & Barker, 2008; Ransome, Renoir, & Hannan, 2012). In this task, mice were exposed to 4 unique objects during a training session, and then exposed to the same 4 objects but with the positions of 2 objects switched (Fig 4A). Mice with normal cognitive function will recognize and spend more time exploring the switched objects. BAC226Q mice, on the other hand, showed no discrimination of the moved objects (Fig 4A) despite spent an equal total amount of time exploring the objects in both training and testing sessions. These results suggest that the BAC226Q mice have impairments in hippocampal-dependent cognitive function at 2 months.

**Figure 4.**
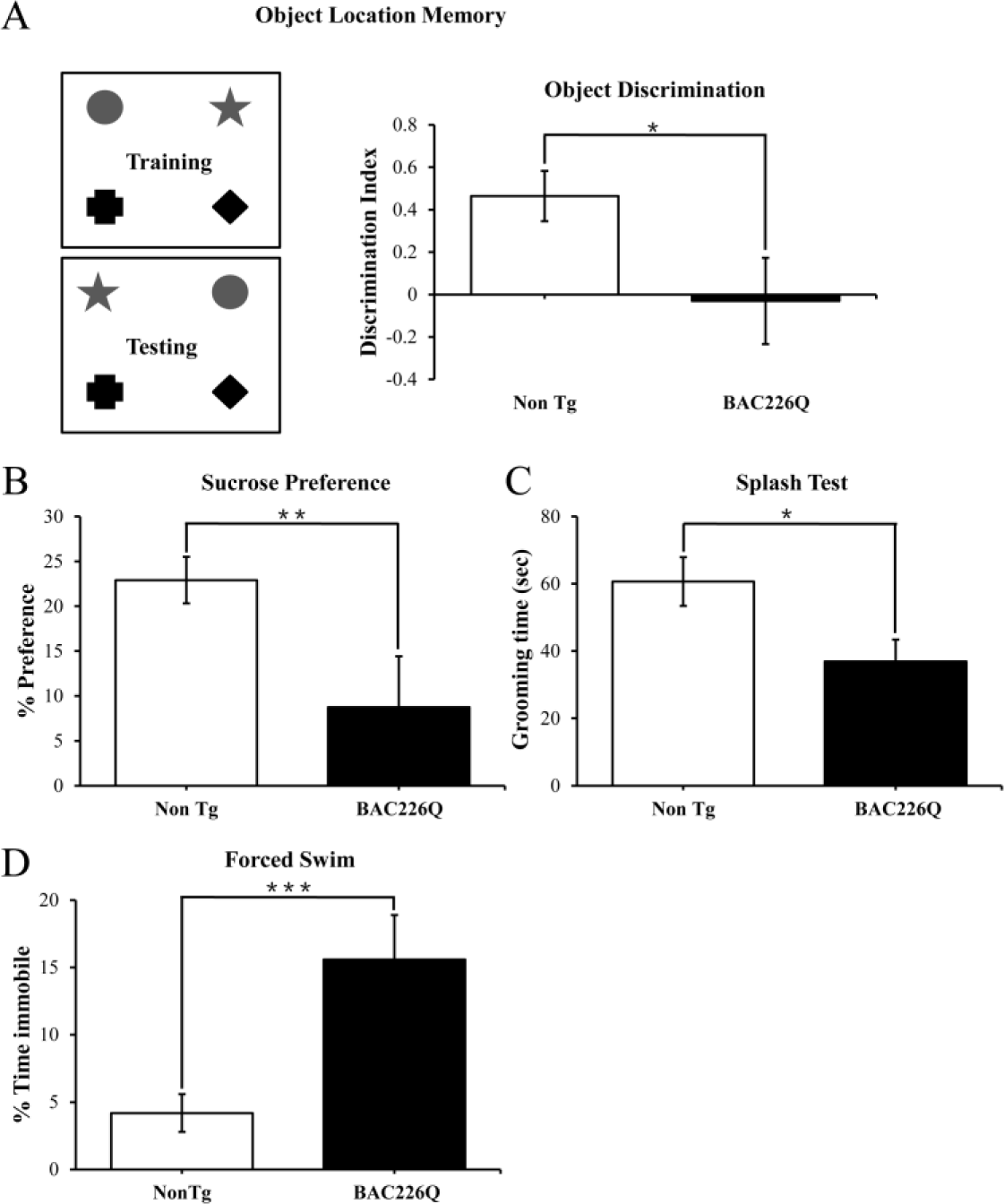
Cognitive and psychiatric disorder-like behavior in BAC226Q mice. All tests used 12 pairs of 2-month-old male BAC226Q mice and non-transgenic sibling controls. BAC226Q mice showed significant decrease in discrimination of moved objects (A, *p* = 0.0485), in sucrose preference (B, *p* = 0.0031), in grooming time in the splash test (C, *p* = 0.035), and in mobile time in the forced swim test (D, *p* = 0.0007). Significance is indicated by *= *p* <0.05, **= *p* <0.01, ***= *p* <0.001.

Depression and anhedonia are also common in HD patients. To test if our mice displayed similar mood phenotypes, we applied three well-characterized behavioral assays. In the sucrose preference task, 2 month old animals were given a choice of regular drinking water or 1% sucrose solution, and consumption of both was measured over 48 hours. While non-transgenic littermates preferred the sucrose solution, BAC226Q mice showed significantly reduced preference (Fig 4B), indicating an anhedonia-like phenotype typical of depression. The splash test is another test of depression-like mood dysfunction in mice. When sprayed with a sticky sucrose solution, BAC226Q mice spent significantly less time grooming compared to non-transgenic littermate (Fig 4C), demonstrating further a depression-like behavior. In the third experiment to explore depression-like phenotypes, we evaluated immobility in the forced swim task. The results were highly significant that BAC226Q mice had three times higher immobility time than non-transgenic littermate controls (Fig 4D). Combined, these three tests indicated a range of depression-like phenotypes including disinterest in rewarding stimuli, apathy and hopelessness in a stressful situation.

### Age-dependent and progressive striatal atrophy and neuronal loss in BAC226Q mice

Post-mortem brains of HD patients have a significant reduction in brain volume, especially in the striatum and deep layer cortex. Age-dependent and progressive striatal atrophy and neuronal loss are cardinal neuropathology of HD patients that are well recapitulated in BAC226Q mice. At 2 months, BAC226Q mice have similar forebrain weight compared to non-transgenic littermates. At 15 months, whole brains from surviving BAC226Q mice weighed 31% less than brains from non-transgenic littermate (Fig 5B). To further evaluate HD-like neuropathology in BAC226Q mice, we used the Cavalieri stereologic estimator to compare striatal volume in 11 month BAC226Q and non-transgenic littermates. The results showed that BAC226Q striatal volume was significantly decreased by 34% compared to controls (Fig A and Fig. 5C). Additionally, similar to the enlarged brain ventricles in HD patients due to brain atrophy, the volume of the ventricles in BAC226Q mice was greatly enlarged by 3.8-fold compared to that of the non-transgenic littermates (Fig 5D). These changes in striatal and ventricular volumes closely mirrored the cardinal pathology in HD patient brains.

**Figure 5.**
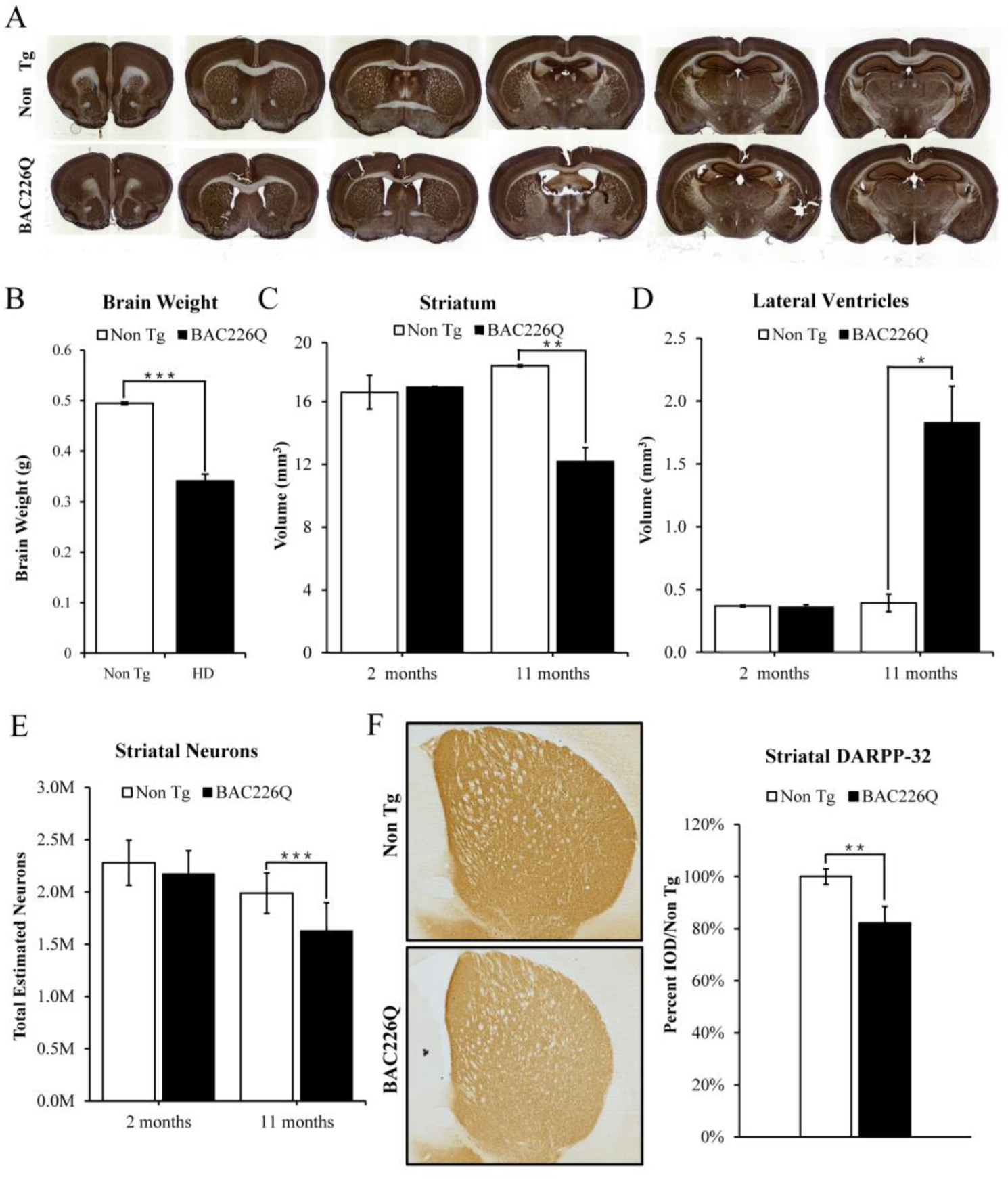
Histopathology of striatal atrophy and neuronal loss in BAC226Q mice. A) Representative micrographs of NeuN stained serial brain sections for 11-month BAC226Q and non-transgenic littermates. B) At 15 months, weights of BAC226Q mice brain were significantly reduced (n=8; *p* = 0.00019). C & D) Striatal atrophy was measured by the Cavalieri stereological estimator. Compared to the non-transgenic littermates, BAC226Q at 2 months had no defects in striatal and ventricle volume. At 11 months, BAC226Q striatal volume was significantly decreased by 34%, n = 4; *p* = 0.0054 (C), and lateral ventricle volume was increased by 379%, p = 0.0127 (D). E) Total striatal neuron count was estimated by an optical dissector stereological estimator. No significant difference was detected at 2 months, but a significant 18% decrease was detected in 11-month BAC226Q (n = 6 BAC226Q, 5 Non Tg, *p* = 0.0002). In every subject counted, the estimated coefficient of error was less than 0.1. F) In 11-month BAC226Q mice striatum, DARPP-32 staining of MSNs was reduced by 18.1% (n=4, *p* =0.0012). In all panels, significance is indicated by ∗ = *p* < 0.05, ∗∗= *p* < 0.01 and ∗∗∗ = *p* < 0.001.

In human HD patients, striatal volume loss is attributed largely to the death of medium spiny neurons (MSNs), which comprise 95% of neurons in the striatum (Oorschot, 1996). To determine whether the striatal volume reduction in BAC226Q mice was caused by neuronal death, we used an unbiased optical fractionator method to count the total striatal neurons in brain tissues from 2 and 11 month animals. At 2 months, no significant differences were detected between genotypes with an average striatal neuron population of 2,280,000±125,000 for non-transgenic and 2,174,000±127,000 for HD mice (Fig 5E). However, at 11 months, non-transgenic littermate control mice had an average of 1,988,000±193,000 striatal neurons, while BAC226Q mice had only 1,631,000±269,000, a difference of 18.0% (Fig 5E). To further evaluate whether the neuronal loss is with MSNs in BAC226Q striatum, MSNs were identified by DARPP-32 immunostaining and quantified by integrated optical density (IOD) in BAC226Q and non-transgenic littermate controls. DARPP-32 IOD was significantly reduced by 18.1% in BAC226Q striatum at 11 months of age (Fig 5F).

### Specific pattern of regional brain atrophy in BAC226Q mice by MRI study

To further characterize the degeneration in the HD mouse brain and to evaluate the feasibility of performing *in vivo* imaging to track disease progression, we imaged 12-month old mice with high-resolution structural MRI and used automated deformation-based morphometry to determine regional brain volumes (Fig 6A). In 12 month transgenic BAC226Q mice, whole brain volume was 373.8±2.95 mm^3^, compared to 522.3±21.7 mm^3^ in non-transgenic controls, a decrease of 28.5% (Fig 6B). In order to make a fair comparison of regional differences between genotypes, we normalized all volumes by whole brain volume to obtain a percent volume for each region before performing statistical tests. We found statistically significant volume changes in several regions of the brain, including the cortex (24.8%, *p*=0.0113, Fig 6C) and striatum (19.4%, *p*=0.0453, Fig 6D). In contrast, there was no significant difference in cerebellar and amygdala volumes between genotypes (Fig 6F and Fig 6G). This is consistent with the observation that the cerebellum and amygdala are largely unaffected in Huntington’s disease patients. Concomitantly, there was a significant increase in the ventricle volume in the BAC226Q mice compared to non-transgenic littermate controls (206.8%, *p*<0.0001, Fig 6E), which was consistent with our Cavalieri stereologic estimator data (Fig 5D). Other regions affected at 12 months included cingulum, stria medullaris, anterior commissure and corpus callosum/external capsule, suggesting a pronounced effect on white matter tracts (Fig 6H). It is important to note that the detection of robust and progressive structural deficits as a biomarker by MRI provides a non-invasive and sensitive method for future therapeutic testing.

**Figure 6.**
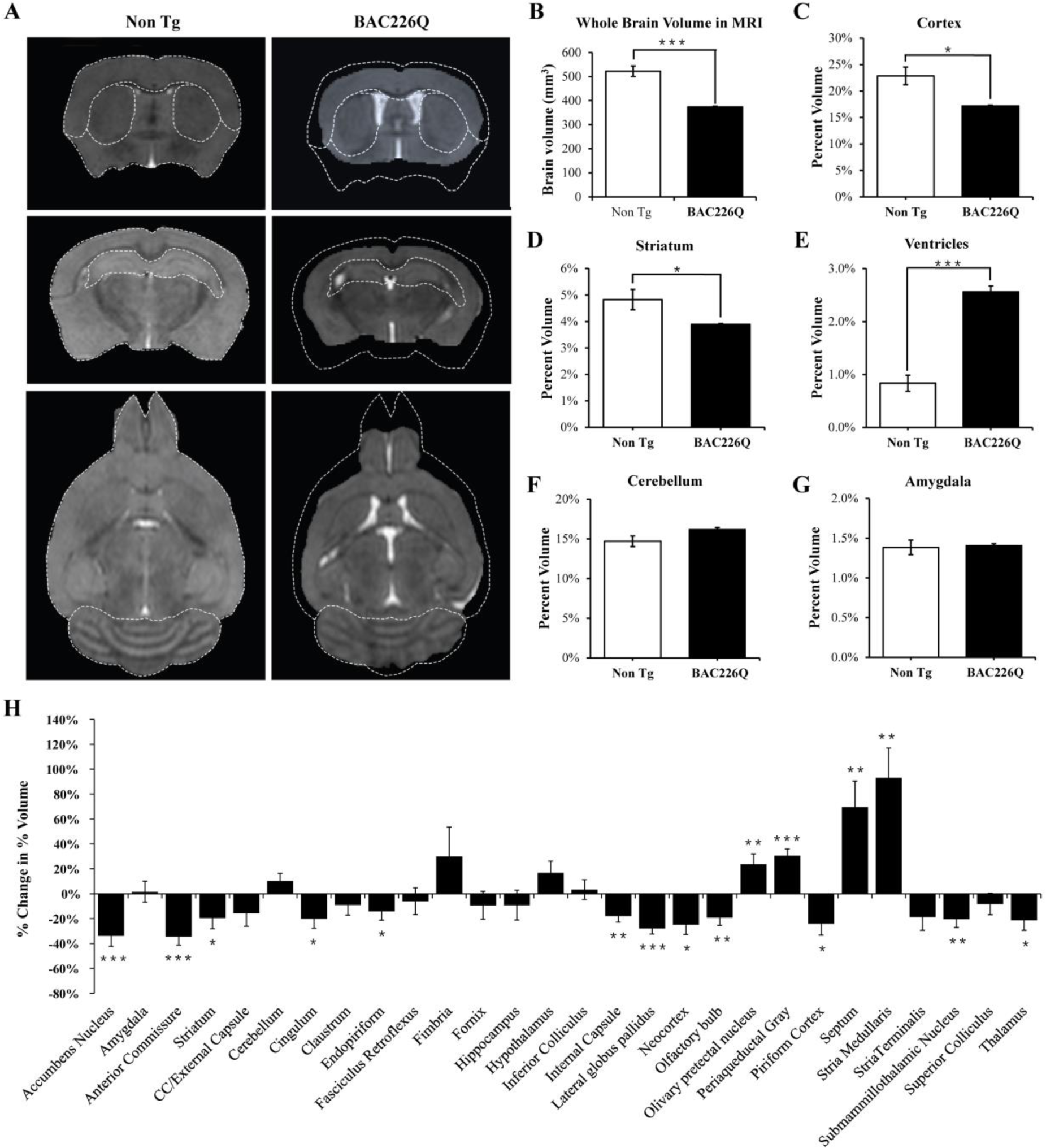
MRI study of regional specific brain atrophy in BAC226Q mice. A) MRI images of 12-month BAC226Q and non-transgenic littermates. B-G) Overall brain volumes were segmented automatically into regional brain volumes by MRI volumetric analysis in BAC226Q mice (n=9) and non-transgenic littermates (n=6). The volumes are presented here as the percentage of whole brain volume. In BAC226Q mice, the decreases of whole brain volumes (28.5%, *p*=0.00022) (B), in relative cortex volume (24.8%, *p*=0.0113) (C) and striatum (19.4%, *p*=0.0453) (D) are significant, as is the increase in ventricle volume (E) (206.8%, *p* <0.0001). There is no significant difference between genotypes in cerebellar (F) and amygdala volume (G). H) Total 28 brain regions were measured by MRI volumetric analysis. The changes of specific brain region volume in BAC226Q mice are presented as the percentage of the same brain area in non-transgenic littermates. In BAC226Q mice, 16 brain regions had significant volume changes (p<0.05). In all panels, significance is indicated by ∗ =*p*< 0.05, ∗∗= *p*< 0.01 and ∗∗∗ =*p*< 0.001

### Aggregate pathology and reactive gliosis in the BAC226Q brain

In post-mortem HD brains, mHtt aggregations are found throughout the central nervous system as an important hallmark (Mangiarini *et al*., 1996). To characterize the distribution patterns of huntingtin aggregations in the specific brain regions of BAC226Q mice, we stained brain tissues from 2-, 4-, and 11-month old BAC226Q and non-transgenic littermate control mice with the S830 antibody raised against N-terminal huntingtin, which specifically recognizes huntingtin aggregates (Fig 7). In agreement with previous findings (Gray *et al*., 2008; Kazantsev, Preisinger, Dranovsky, Goldgaber, & Housman, 1999; Li, Li, Yu, Shelbourne, & Li, 2001), two types of huntingtin aggregates were observed in BAC226Q mice, nuclear inclusions (NIs) and neuropil aggregates (NAs).

**Figure 7.**
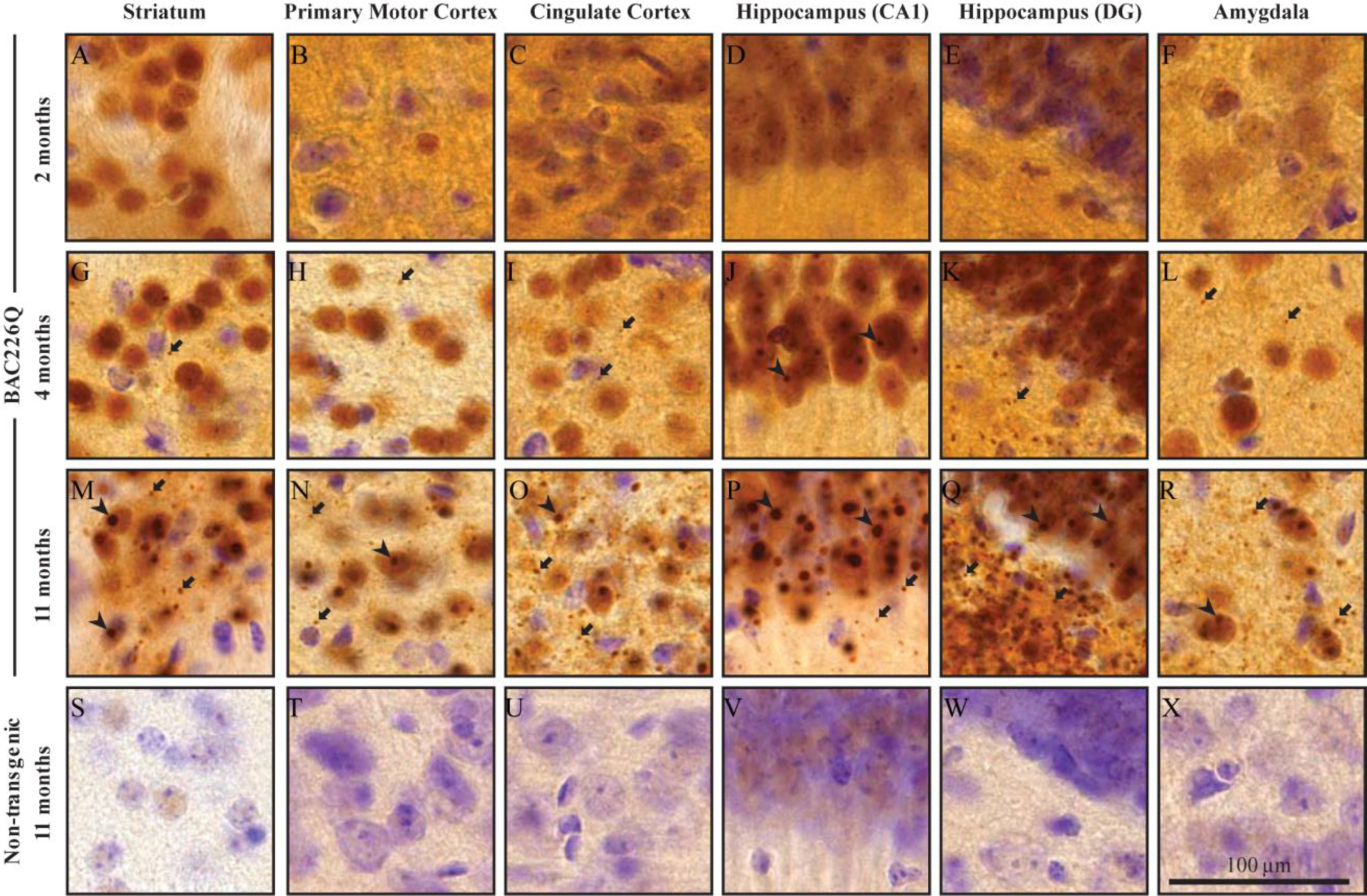
Widespread and progressive mHtt aggregate pathology in BAC226Q brain. Huntingtin aggregates were stained by the S830 antibody. In BAC226Q, mHtt aggregates were undetectable at 2 months (A-F), detected as small aggregates (arrows) and nuclear inclusions (arrowheads) at 4 months (G-L), finally developed to large and numerous cytosolic aggregates and nuclear inclusions at 11 months (M-R). In contrast, no aggregates were detected in non-transgenic littermates (S-X). Regions shown are striatum (A, G, M, S), primary motor cortex (B, H, N, T), cingulate cortex (C, I, O, U), CA1 of hippocamus (D, J, P, V), dentate gyrus (E, K, Q, W) and amygdala (F, L, R, X).

At 2 months, neuron cell bodies were diffusely stained by the S830 antibody throughout the brain in BAC226Q mice (Fig 7A-F), but no immunoreactivity detected in non-transgenic littermates (Fig 7S-X). This suggests that at 2 months only soluble mHtt protein is present. In contrast, punctate staining of mHtt protein by S830 became evident in several regions of the brain at 4 months, including striatum (Fig 7G), motor (Fig 7H) and cingulate cortex (Fig 7I), hippocampus (Fig 7J, K) and amygdala (Fig 7L) in BAC226Q mice, but not control mice. At 11 months, aggregate pathology was much more severe in BAC226Q mice (Fig 7M-R).

Reactive gliosis is another cardinal pathology of HD in addition to selective neuronal loss. We analyzed reactive gliosis by immunohistochemical detection of GFAP, the astrocyte-specific marker, in BAC226Q mice and non-transgenic littermate. An 80% increase in reactive gliosis was found in deep layer cortex and striatum at 11 month BAC226Q mouse brain (Fig 8).

**Figure 8.**
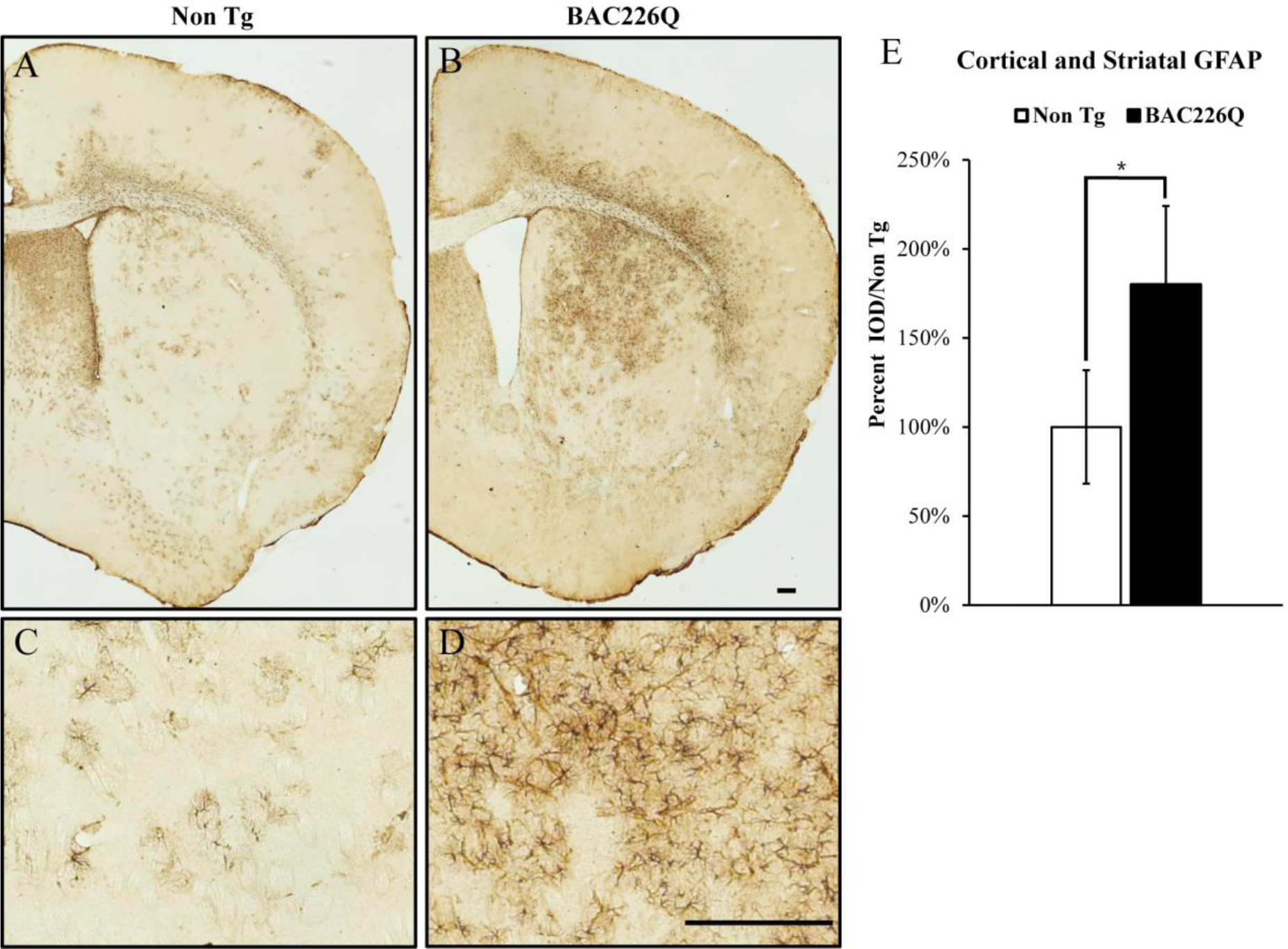
Gliosis in striatum and deep cortical layers of BAC226Q brain. Reactive astrogliosis is analyzed by immunohistochemical staining with GFAP antibody in 11-month BAC226Q and non-transgenic littermate brains. Gliosis is prominent in BAC226Q (B&D), but not in control littermate brains (A&C) (n=4, *p* =0.032).

## Discussion

Many mouse models have been developed for investigating the underlying pathogenic mechanisms and testing therapeutic methods. Categorically, a mouse model will be compelling if it meets two criteria simultaneously: accurate recapitulation of cardinal HD phenotypes in one mouse model rather than separately in several models, and absence of erroneous and unwanted false phenotypes that do not correlate with human HD. It should be emphasized that the novel BAC226Q reported here is such an example.

### Robust and faithful recapitulation of HD

As presented in the Results section and Table 2, the BAC226Q mouse model has validity and fidelity in a full spectrum from genomic DNA, protein, subcellular/cellular pathology, histopathology, specific brain area atrophy, cognitive and psychiatric disorder-like phenotypes, motor behavioral deficits, weight loss and shortened life spans.

**Table 1.** Recapitulation of cardinal HD phenotypes in BAC226Q mice.

|  | HD Patients | BAC226Q Mice |
| --- | --- | --- |
| Neuropathology | Brain atrophy | Yes |
|  | Neuron loss | Yes |
|  | mHTT aggregations | Yes |
|  | Reactive gliosis | Yes |
| Progressive motor deficits | Bradykinesia | Yes |
|  | Incoordination | Yes |
|  | Chorea | Yes |
|  | Dystonia | Yes |
| Non-motor phenotypes | Psychiatric symptoms | Yes |
|  | Cognitive defects | Yes |
| Reduced life span |  | Yes |
| Weight loss |  | Yes |

At the genomic level, the human BAC clone contains the complete human HTT gene with its endogenous intron-exon structures and 5’, 3’ regulatory regions. This is important for testing future gene editing approach such as CRISPR/Cas9 mediated targeting. BAC226Q transgenic mice express full length human HTT with 226 CAG-CAA repeats, which yielded robust and early on-set HD phenotypes. It should be noted that 226Q is within the range of mutations identified in human patients (Nance *et al*., 1999). In additional consideration for practical usage, it is worth noting that the CAG-CAA mixture gives BAC226Q mice an important advantage of complete polyQ length stability between generations and individuals.

Importantly, BAC226Q mice do not have spurious phenotypes that sometimes exist in other models. For example, unlike other models with severe developmental problems, BAC226Q mice develop normally and subsequently have age-dependent and progressive neurodegeneration. Also, unlike other models that showed significant body weight gain which is opposite to human HD, BAC226Q mice have progressive weight loss and reduced lifespan, consistent with human HD weight loss as a hallmark (Gaba *et al*., 2005; Stoy & McKay, 2000). Interestingly, since weight loss is also observed in BAC-225Q model with full-length mouse mutant HTT (Van Raamsdonk *et al*., 2006; Wegrzynowicz *et al*., 2015), it seems that weight loss phenotype is polyQ-length dependent rather than human or mouse HTT species dependent.

Compared to other human full length HTT models, a significant advantage of BAC226Q mice is its much earlier onset, much more robust and faster progressing phenotypes including early hyperkinetic / late hypokinetic biphasic motor dysfunction. HD patients develop hyperkinetic chorea first and as the disease progresses, hypokinesia and dystonia subsequently. Parallel to the patients motor symptom progression, BAC226Q mice have severe hyperkinesia including involuntary choreiform movement by 12-16 weeks and show reduced mobility after 10 months.

An important but less well-studied aspect of HD is psychiatric and cognitive impairments, which often occur decades before the onset of motor symptoms in HD mutation carriers. In addition to motor dysfunctions, BAC226Q mice show non-motor phenotypes at 2 months, before the onset of motor deficit and neuropathology, which is the same temporal sequence as in patients. Thus BAC226Q is an appropriate and powerful tool to study the mechanisms underlying psychiatric and cognitive deficits.

### A model well suited for preclinical investigations

Several features make BAC226Q well suited for preclinical studies. First, BAC226Q accurately recapitulates HD phenotypes. Therefore, candidate drugs or approaches for disease modification can be tested for their abilities to rescue these highly relevant phenotypes at multiple levels. Second, BAC226Q has normal development and very early onset of phenotypes that are progressive and robust. This provides a long window of observation for testing the efficacy of therapeutic candidates. Third, the remarkable region-specific brain atrophy revealed by high-resolution structural MRI can be used as a biomarker and readout in preclinical studies longitudinally without sacrificing the animals. Fourth, the CAG-CAA mix makes the polyQ length stable between generations and among individuals, which is advantageous in reducing individual variability. Last but not least, BAC226Q has human genomic DNA as the transgene, which is very appropriate testing gene-editing such as CRISPR/Cas9 strategy in preclinical studies.

### Insights of mHtt toxicity in BAC226Q

Although we only conducted an initial analysis of BAC226Q mice, we have already generated data that can be used to clarify some of the questions in the field.

Huntingtin aggregates were first identified in R6 mice expressing exon 1 of mutant huntingtin and subsequently identified in human tissue (Mangiarini *et al*., 1996). It is widely postulated that Htt aggregates are the source of HD toxicity. Since N-terminal huntingtin is much more susceptible to aggregation compared to the full length protein (Ratovitski *et al*., 2009), and causes a more severe disease phenotype in mice, there has been a hypothesis that full length mHtt need to be cleaved into fragments to exert toxicity. Our data do not support this hypothesis, because in the life span of BAC226Q mice, fragmented mHtt is hardly detectable. It seems that in BAC226Q, full length mHtt is toxic and sufficient to drive pathogenesis.

Another important question is whether mHtt has a dominant gain-of-toxic function, or a combination of loss of function of the wild type allele. In BAC226Q, mHTT was not overexpressed, and the two alleles of the wild type HTT exist. The fact that BAC226Q mice developed such robust HD-like phenotype seems to lend support to the “gain-of-function” hypothesis. The direct implication is that deleting mutant HTT alleles will be sufficient to benefit patients in CRISPR/Cas9 mediated gene-targeting as a therapeutic method.

## Conclusions

In this study, we report the generation and analyses of a novel BAC226Q mouse, which accurately recapitulates the cardinal HD phenotypes including body weight loss, HD-like characteristic motor behavioral impairment, cognitive and psychiatric symptoms, and classic HD neuropathology changes such as significant neuronal death in striatum and cortex, widespread mHtt aggregation pathology and reactive gliosis. Therefore, this model will be valuable for mechanistic studies and therapeutic development of HD, especially for the preclinical genetic therapies targeting human mHTT.

## Materials and methods

### BAC engineering and generation of transgenic mice

A BAC containing the full-length wild-type human HTT gene was modified to express a full-length HTT with a stable, expanded polyglutamine tract. A plasmid containing exon1 of HTT with 226 mixed CAG-CAA repeats was a gift from Dr. Alex Kazantsev (MIT). The sequence corresponding to 226Q was inserted into the HTT gene by homologous recombination using our standard protocols (Gong, Yang, Li, & Heintz, 2002). Fingerprinting analysis by genomic Southern blots detected no unwanted rearrangements or deletions in the modified Htt-226Q BAC. The modified BAC was sequenced to confirm that there were no unwanted mutations other than the intended 226Q insertion. Finally, the full-length human Htt-226Q BAC copy number and the insertion site in mouse genome were determined by whole genome sequencing and bioinformatics analysis, performed by Novogene Corporation.

### Animal husbandry

Mice were housed in a temperature and humidity controlled specific pathogen-free (SPF) facility under a 12-hour light/dark cycle schedule with food and water available *ad libitum*. Transgenic mice were bred with wild-type FVB/N mice (Taconic). It is important to note that BAC226Q in the Taconic FVB/N mouse background exhibited the most robust and consistent phenotypes. Genotyping was determined by PCR of genomic DNA isolated from tail snips. Genotyping primers are located in intron 2 and are specific to human huntingtin (forward primer: 5’-GTA TAT GCT GCT GCC TGC AA-3’; reverse primer: 5’-AGG GGA CAG TGT TGG TCA AG-3’), producing a 403bp PCR fragment. Non-transgenic littermates were used as controls. All experimental protocols were approved by the Weill Cornell Medicine and Peking University animal care and use committee.

### Behavioral study

#### Open Field Test

Mice were monitored individually with the VersaMax Animal Activity Monitoring System (Accuscan Instruments). The activity monitor consists of a 16x16 grid of infra-red beams at floor level to track horizontal position with 16 additional beams at a height of 3” to detect vertical activity. Horizontal and vertical activities were measured by beam breaks recorded over a period of 1 hour using the VersaMax software.

#### Cylinder Test

The cylinder test was performed during the dark phase of the light-dark cycle. Each animal was placed in a clear acrylic cylinder and recorded on video for 5 minutes. A mirror was placed below the cylinder to provide a view of the animal from below. Videos were analyzed using ImageJ.

#### Rotarod

8-week-old mice were trained in three 60-second trials at a constant speed of 10 rpm. During the training trials, mice which fell were gently returned to the rotarod. Starting at 9 weeks, mice were tested once a week on an accelerating rotarod (5-45 rpm over 5 minutes) in 3 trials, with about 15 minutes rest between trials.

#### Sucrose preference test

Individually housed mice were trained to drink from two identical water bottles on day 0. On day 1, one water bottle was replaced with a 1% sucrose solution and on day 2, the positions of bottles were switched. The volumes of plain water and sucrose solution before and after experiments were recorded. % preference was calculated by the following formula 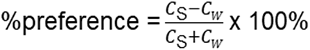, where CS and CW are sucrose consumed and water consumed, respectively.

#### Splash test

Mice were sprayed on their dorsal coat with a 10% sucrose solution, placed in an empty cage, and recorded for 5 minutes. Duration of grooming activity was scored by a person blind to genotype.

#### Forced swim task

Mice were placed individually into a cylinder of room temperature water (25±1°C) and recorded for 5 minutes. Immobility (time spent floating and not swimming) was scored manually by a person blind to genotype.

#### Object location memory

Mice were introduced to an open field chamber for five minutes for 3 days. On day 4, four unique objects were placed to the open field chamber. On day 5, mice were presented with the same four objects but the position of two objects was switched. Discrimination Index (DI) was calculated using the formula 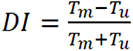 where *T_m_* and *T_u_* are time spent exploring the moved and unmoved objects, respectively.

### Western blot

Freshly dissected mouse brains were homogenized by a Precellys tissue homogenizer in ice-cold RIPA buffer supplemented with Complete Protease Inhibitor Cocktail tables (Roche). The supernatant was collected after centrifugation in a refrigerated centrifuge (4°C) at 16,000 xg for 10 minutes. Protein concentration was determined by a BCA Assay (Pierce), and an equal amount of protein was loaded in each well of NuPAGE 4-15% Tris-Acetate gels or 15 x 22 cm, 8% polyacrylamide gels for SDS-PAGE. Proteins were transferred to a PVDF membrane (Immobilon-FL) and probed with anti-Huntingtin 1C2 or S830. An odyssey infra-red imager was used to visualize the blots (LI-COR biosciences)

### Neuropathology

#### Tissue preparation

Mice were anesthetized with a ketamine/xylazine cocktail and transcardially perfused with PBS followed by 4% freshly prepared, ice-cold paraformaldehyde (PFA). Brains were removed and post fixed for 24 h in PFA, then transferred into PBS for storage. Brains were mounted in 4% agarose and coronal sections were obtained with a vibratome (40um and 60um thickness). Serial sections were collected into 10 wells so that each well contained every 10th section. Floating sections were stored in anti-freeze buffer (30% glycerol and 30% ethylene glycol in PBS) at -20°C.

#### Immunostaining

Floating sections were stained with the NeuN antibody (Millipore) at a 1:1000 dilution, the S830 anti-Htt antibody (a gift from Dr. Gillian Bates) at a 1:25,000 dilution, the anti-GFAP antibody (Abcam) at a 1:2000 and the DARPP-32 antibody (Abcam) at a 1:7500 for overnight at 4°C. Subsequent incubations in secondary antibody, ABC solution, and color development were performed according to instructions provided with the ABC and DAB substrate kits (Vector Labs). Brain sections stained by NeuN antibody and S830 anti-Htt antibody were counter-stained with cresyl violet.

#### Stereology

Stereologic measurements were obtained by Stereo Investigator (MBF Biosciences) with a Zeiss Axiophot2 microscope. The optical fractionator probe was used to estimate total striatal neuron number in every 10th section. The counting frame was 30 x 30s in the NeuN antibody stained brain sections with an optical dissector depth of 8 or 18µm for 40 and 60 micron cut sections, respectively with 2µm guard zones. At least 300 neurons were counted for each animal, and the estimated coefficient of error in each case was less than 0.1. Striatal and ventricle volumes were determined using the Cavalieri estimator on the same slides with a grid size of 100 x 100 microns oriented at a random angle.

#### MRI volummetry study

MRI scanning and computational analysis were performed as previously described (Cheng *et al*., 2011). Female BAC226Q and wild-type FVB mice aged 12 months were scanned by a 9.4 Tesla MR scanner with a triple-axis gradient and animal imaging probe. The scanning was performed *in vivo* under isofluorane anesthesia. The resulting images were aligned to a template by automatic registration software and signal values were normalized to ensure consistent intensity histograms. A computer cluster running Large Deformation Diffeomorphic

Metric Mapping (LDDMM) was used to automatically construct non-linear transformations to match anatomical features and perform automatic segmentation of specific brain regions. The volume of each segmented structure was normalized by total brain volume and analyzed for each genotype.

### Statistics

Data were showed as mean± SEM (standard error of the mean) unless otherwise noted. Student’s t-test (unpaired) or one-way ANOVA were used to compare transgenic and control groups in each experiment with significance indicated by *p*-values less than 0.05. *p*-values, SEM, means and standard deviations were calculated with Microsoft Excel 2010 and Prism software. Survival curves were compared with the log-rank (Mantel-Cox) test.

## Acknowledgements

This project was partially supported by Hereditary Disease Foundation, Weill Cornell Medical College and Peking University School of Life Sciences. The authors declare no competing financial interests.

## Supplementary Material

Video 1. Chorea-like movement in BAC226Q mice at 14 weeks

Video 2. Rapid circling behavior in BAC226Q mice at 16 weeks

Video 3. Cylinder test in BAC226Q mice at 16 weeks

## Article and author information

Chenjian Li,

- School of Life Sciences, The MOE Key Laboratory of Cell Proliferation and Differentiation, Peking University, Beijing, China
- Former Affiliation: Department of Neuroscience, Weill Cornell Graduate School of Medical Sciences, New York, NY, USA

Present address:

School of Life Sciences Building Room # 630, Peking University, Beijing, 100871, China Phone: 86-10-6275-6459

## Contribution

Conception and design, Acquisition of data, Analysis and interpretation of data, Supervision, Funding acquisition, Investigation, Methodology, Writing - original draft, Project administration, Writing - review and editing

## Competing interests

No competing interests declared.

Sushuang Zheng

School of Life Sciences, The MOE Key Laboratory of Cell Proliferation and Differentiation, Peking University, Beijing, China

**Present address:**

School of Life Sciences Building Room # 630, Peking University, Beijing, 100871, China **Contributio**n: Conception and design, Acquisition of data, Analysis and interpretation of data, Writing - original draft, Writing - review and editing

**Competing interests:** No competing interests declared.

Sushila A Shenoy

Department of Neuroscience, Weill Cornell Graduate School of Medical Sciences, New York, NY 10021, USA

**Contribution**: Conception and design, Acquisition of data, Analysis and interpretation of data, and Drafting or revising the article

**Competing interests:** No competing interests declared.

Wencheng Liu

Department of Neuroscience, Weill Cornell Graduate School of Medical Sciences, New York, NY 10021, USA

**Contribution**: Acquisition of data, Analysis and interpretation of data, and Writing - review and editing

**Competing interests:** No competing interests declared.

Yuanyi Dai

School of Life Sciences, Peking University, Beijing, China

**Contribution**: Acquisition of data, Analysis and interpretation of data, and Writing - review and editing Competing interests: No competing interests declared.

Yuanxiu Liu

School of Life Sciences, Peking University, Beijing, China

**Contribution**: Acquisition of data, Analysis and interpretation of data and Writing - review and editing

**Competing interests:** No competing interests declared.

Zhipeng Hou

The Russell H. Morgan Department of Radiology and Radiological Sciences, Johns Hopkins University School of Medicine, Baltimore, Maryland, USA.

**Contribution**: acquisition of data, analysis and interpretation of data, and revising the article

**Competing interests:** No competing interests declared.

Susumu Mori

The Russell H. Morgan Department of Radiology and Radiological Sciences, Johns Hopkins University School of Medicine, Baltimore, Maryland, USA.

**Contribution**: acquisition of data, analysis and interpretation of data, and revising the article

**Competing interests:** No competing interests declared.

Wenzhen Duan

Translational Neurobiology Laboratory, Baltimore Huntington’s Disease Center, Johns Hopkins University School of Medicine, Baltimore, Maryland, USA.

**Contribution**: Acquisition of data, Analysis and interpretation of data, Methodology, and Writing - review and editing

**Competing interests:** No competing interests declared.

